# eDNA metabarcoding and whole genome sequencing detect European-American Eel hybrids in northeastern Canada

**DOI:** 10.1101/2025.04.21.649106

**Authors:** Samantha E. Crowley, Tony Kess, Paul Bentzen, Victoria Neville, Chelsea Bloom, Nicole Smith, Nicole Fahner, Lesley Berghuis, Kerry Hobrecker, Mehrdad Hajibabaei, Steven J. Duffy, Ian R. Bradbury

## Abstract

American and European Eels (*Anguilla rostrata* and *Anguilla anguilla*, respectively) are sister species of conservation concern, as they have declined in abundance in recent decades. Although there is evidence of gene flow between the two species, most previous hybrid detections have been found in Iceland. Here we expand upon existing reports of interspecific hybrids using a combination of an environmental DNA (eDNA) survey and whole genome sequencing of 347 eels from Newfoundland and Labrador. Metabarcoding of eDNA samples at multiple loci detected the presence of European Eel mitochondrial DNA (mtDNA) at eight locations primarily along the south coast of Newfoundland. Genome sequencing of eels collected in the region revealed three hybrid individuals with levels of European admixture consistent with one first generation hybrid and two North American backcrosses, all with European mtDNA. This work supports the presence of a previously unknown, seemingly localized region of hybrid occurrence in southern Newfoundland and expands our understanding of the relationship between these species across the North Atlantic.

## Introduction

American Eel (*Anguilla rostrata*) and European Eel (*Anguilla anguilla*) represent two sister species occurring on respective sides of the Atlantic Ocean (Tagliavini et al. 1996; Aoyama et al. 2001; Lin et al. 2001; Aoyama 2003) and have been fished by people for millennia (i.e. Dekker 2019). As long-lived, semelparous, facultatively catadromous species, *A. rostrata* and *A. anguilla* spend the majority of their lives in freshwater and coastal habitats. Both species have shown marked declines over the past several decades, attributed to factors such as overfishing, habitat degradation and obstruction (i.e. dams), introduced parasites, pollution, and climate change (COSEWIC 2012; ICES 2023). The commercial catch of *A. rostrata* is estimated to have fallen to 50% of its pre-1980 level in certain regions, and the recruitment of juvenile *A. anguilla* has declined by an estimated 90-99% (reviewed by Drouineau et al. 2018). *A. rostrata* and *A. anguilla* are listed under the IUCN Redlist (https://www.iucnredlist.org), as “endangered” and “critically endangered” respectively. Conservation of both species is impeded by knowledge gaps surrounding many aspects of their biology, ecology, and life history.

On the whole, recent work supports the case of true panmixia within each species (Als et al., 2011; Côté et al., 2013; Enbody et al., 2021; Gagnaire et al., 2012; Palm et al., 2009; J. M. Pujolar et al., 2014; Ulmo-Diaz et al., 2023) with significant divergence between species, yet the mechanisms underpinning genetic and geographic isolation of the two species remain largely unknown. Genetically controlled differences in developmental timing are thought to play a role in their isolation (e.g. Bernatchez et al. 2011; Jacobsen et al. 2014), since *A. anguilla* leptocephali tend to drift much further and longer than *A. rostrata* leptocephali. Hybrids have been observed, mainly in Iceland (e.g. Avise et al. 1990; Albert et al. 2006; Gagnaire et al. 2009), though limited admixture has been reported in each species in their continental ranges (Albert et al. 2006; Wielgoss et al. 2014). To this day, no observations of F1 hybrids have been validated outside of Iceland (to our knowledge). However, recent work suggests the potential presence of hybrid *A. anguilla/A. rostrata* outside of Iceland (Crowley et al. 2024), opening the door for new investigation into the relationship between these two species. Ultimately, the spatial scale of hybridization as well as the broader genomic consequences remain poorly understood and necessitates further evaluation.

In this study, we investigated the potential presence of European-American Eel hybrids in southern Newfoundland, Canada, and addressed the following objectives. First, we investigated the presence and robustness of European Eel environmental DNA (eDNA) detections in watersheds around Newfoundland and Labrador using a recently published multi-locus eDNA metabarcoding dataset from 2019-2021 (Crowley et al. 2024). Second, we used low-coverage whole genome sequencing of eel tissue collected in 2020-2022 to explore the presence of European Eels or their offspring in the region. Finally, we used these genomic data to investigate the direction of potential hybridization and to quantify the level of European Eel admixture in hybrid individuals detected. This study builds on previous detections of European/American Eel hybrids in Iceland (Avise et al. 1990; i.e. Albert et al. 2006) and expands our understanding of the frequency and distribution of hybrids of these two species across the North Atlantic.

## Methods

### eDNA Field Sampling and Data Processing

We sampled 107 coastal river sites across the island of Newfoundland in 2020 and 67 sites in Labrador in 2019 and 2021 for eDNA (Figure 1A); see Crowley et al., (2024) for full details of field and lab sampling, and bioinformatic processing. Samples were taken in triplicate (in addition to a field blank) at each site and filtered through 0.45 µm membranes with an eDNA sampler (Smith-Root). DNA was extracted using a DNeasy PowerWater kit (Qiagen, Hilden, Germany). Three mitochondrial DNA (mtDNA) markers were selected for metabarcoding (12S ribosomal RNA region: 12S Teleo (Valentini et al. 2016), 12S MifishU (Miya et al. 2015); and cytochrome oxidase subunit 1 region: CO1 MiniFISH-E (Shokralla et al. 2015)), amplified in triplicate PCRs, and pooled. Amplicons were indexed using unique dual indices, randomized across sampling years into four batches, and sequenced on the NovaSeq 6000 (Illumina, San Diego, USA) using v2 SP and S1 300 cycle sequencing kits, with a per-sample, per-marker target depth of 2 million reads. Raw reads were subsequently processed using a custom pipeline (Creedy et al. 2023) (code available for this pipeline at https://github.com/samcrow93/NL_eDNA/tree/main/Scripts/NL_eDNA_scripts). Briefly, adapters, indices, and primer sequences were removed using cutadapt v2.10 (Martin 2011), forward and reverse reads merged using PEAR v0.9.11 (Zhang et al. 2014), quality, length, and chimaera filtered using VSEARCH v2.21.1 (Rognes et al. 2016), and taxonomy assigned using the blast+ tool (v2.13.0) against the NCBI nucleotide database.

**Figure 1.**
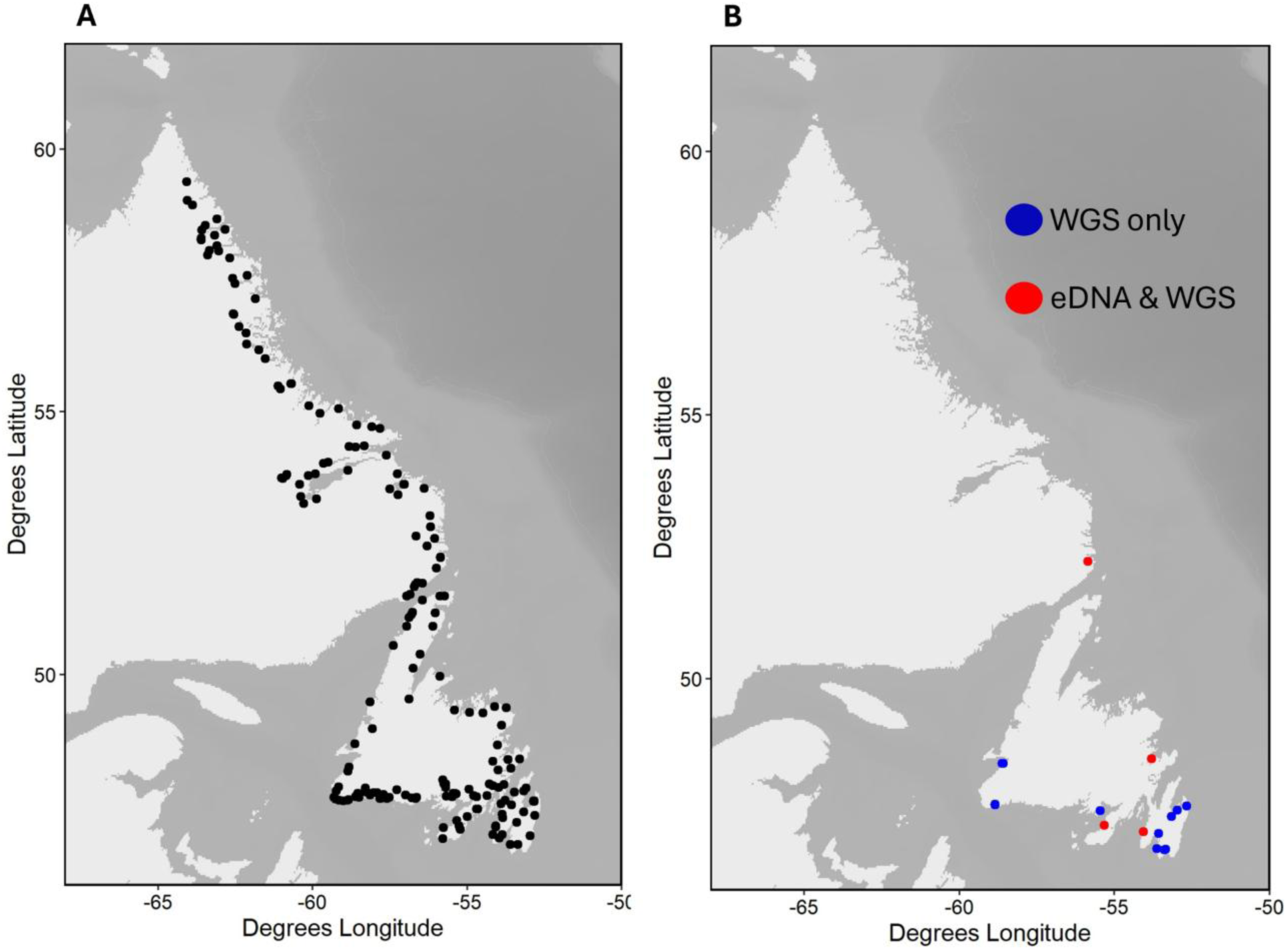
A) Locations of eDNA water sampling in 2019-2021; and B) locations of sampling for eels for whole genome sequencing in 2020-2022, with sites sampled using both eDNA and whole genome sequencing (WGS) shown in red and sites sampled only for WGS shown in blue.

### Low-Coverage Whole Genome Field Sampling and Sequencing

We sampled eels from 14 locations in southern Newfoundland in 2022 (one of which was also sampled in 2021), and from a single location in Labrador (St. Charles River) in 2020 and 2022 (Figure 1b). Four of the southern Newfoundland locations had been sampled during the 2020 eDNA survey. A total of 347 eels were collected by electrofishing (Smith-Root LR-24 Electrofisher), eel traps, and fyke nets; see Table 1 for full sampling details for each site. Fin clips were taken from the pectoral fin and stored in 95% ethanol for later DNA extraction. Specimens were handled following Canadian Council on Animal Care (CCAC) guidelines, and samples taken via electrofishing were collected under an Experimental License for the Salmonids Section of Fisheries and Oceans Canada Newfoundland Region (NL-6664-22), while those collected with fyke nets and traps were in conjunction with individual licensed harvesters and thus exempt from additional permitting after assessment by the Fisheries and Oceans Animal Care Committee.

**Table 1.**
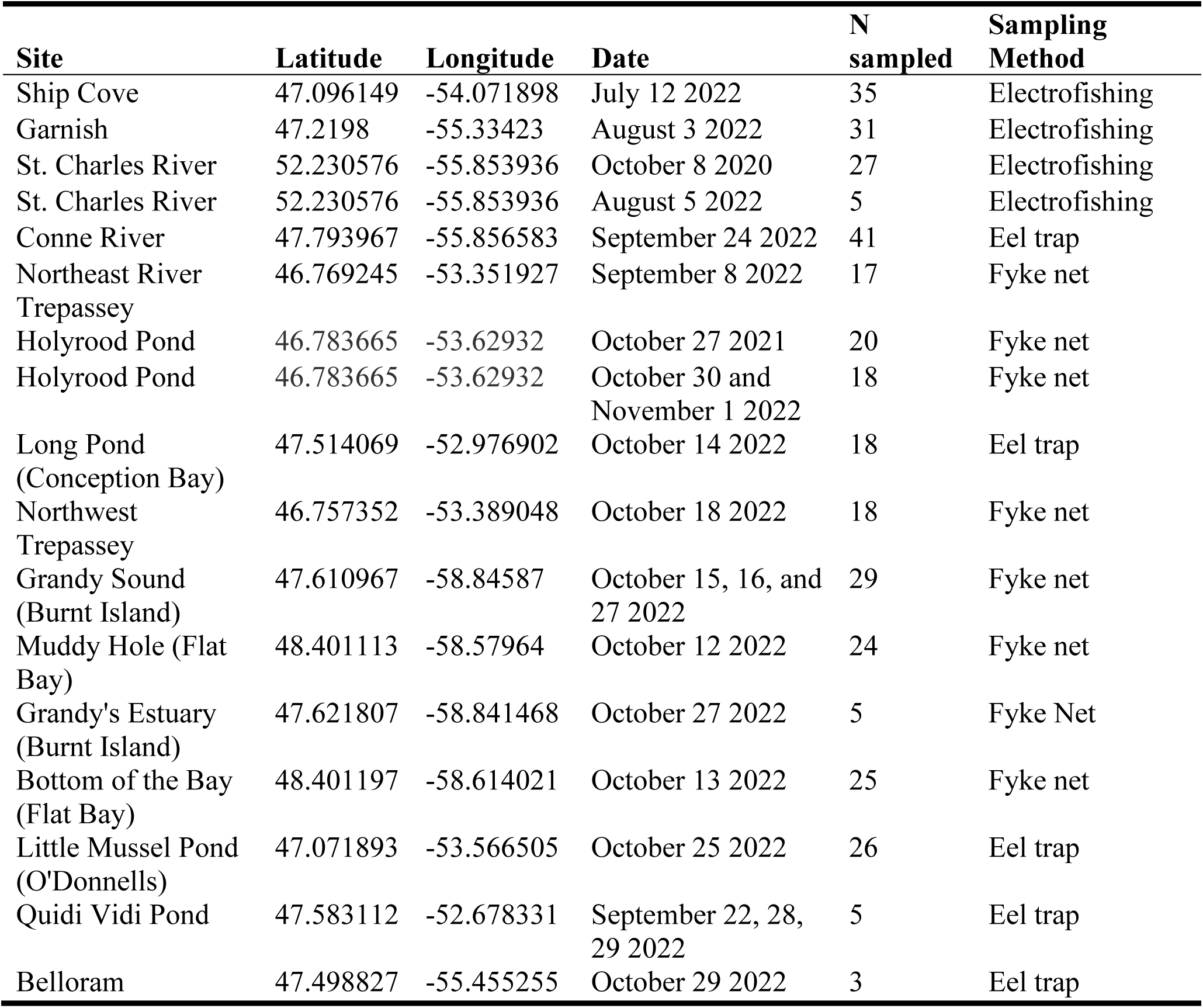
Details of eels sampled at sites across Newfoundland and Labrador from 2020-2022.

We conducted low-coverage whole genome sequencing (LC-WGS) of eels sampled across these 15 sites. Genomic DNA was extracted using DNeasy 96 Blood and Tissue kit (Qiagen) following the manufacturer’s protocol. The quality of the extracted DNA was checked on a 1% agarose gel and quantity was measured using Qubit HS dsDNA quantification kit (ThermoFisher). Following normalization, sequence-ready DNA libraries were prepared using DNA Prep kit (Illumina). Individual libraries were pooled, and library size was quantified on a 2100 Bioanalyzer (Agilent Technologies). Libraries were sequenced on an Illumina NovaSeq6000 S4 at The Genome Quebec Centre d’Expertise et de Services.

### Bioinformatic Processing of Nuclear Data

In addition to the 347 samples collected in southern Newfoundland and Labrador (NL), we also downloaded raw low-coverage whole genome sequence data for 493 eel samples previously published by Enbody et al., (2021), the majority of which were sampled in Europe and identified via genomic analyses as *A. anguilla*, with a smaller number sampled in North America and identified as *A. rostrata*. The main purpose of incorporating these additional samples in our analyses was to be able to compare genomic variation from our eels sampled in NL to genomic variation present in known *A. anguilla* samples. The bioinformatic processing of all raw sequencing data (as well as all downstream analyses) was performed using code available at https://github.com/samcrow93/Two-Eel-or-not-two-Eel. Briefly, we used fastp v0.23.1 (Chen et al. 2018) to remove adapter content from raw reads, using default parameters. The trimmed reads were subsequently aligned to the newest version of the *A. anguilla* reference genome, assembled by the Vertebrate Genomes Project, fAngAng1.pri (NCBI accession: GCF_013347855.1) using bwamem2 v2.2.1 (Li 2013). We then used the Genome Analysis Toolkit (GATK v4.4.0.0, (Van der Auwera and O’Connor 2020)) to remove duplicated reads (function MarkDuplicates) and realign reads around potential insertions and deletions (v3.7; functions RealignerTargetCreator and IndelRealigner).

We used the program ANGSD v0.940 (Analysis of Next Generation Sequencing Data; (Korneliussen et al. 2014) to estimate genotype likelihoods using the SAMtools method and output in Beagle format (-gl 1, doGLF 2), with allele frequencies subsequently estimated from genotype likelihoods using fixed major and minor alleles (-doMajorMinor 1 and -domaf 1), and posterior probabilities of the genotypes calculated by using the frequencies as a prior (-dopost 1). During these calculations, individual bases with quality score below 30 were discarded (minQ 30). Additionally, reads whose mates could not be mapped, reads that did not map uniquely, and reads with mapping quality below 30 were also discarded (-only_proper_pairs 1, -minmapQ 30, - uniqueOnly 1, -remove_bads 1). Sites were discarded if fewer than ∼80% of individuals had data (-minInd 670), or if total sequencing depth across all individuals at that site was below 1700 (equal to approximately two times the total number of individuals; -setMinDepth 1700). We used an estimated minor allele frequency cutoff of 0.05 (-minMaf 0.05). For sites passing these filtering criteria, depth was calculated using -doCounts 1 and -dumpCounts 2. Average coverage for each of the 19 chromosomes (plus the mitochondrial genome) was calculated on a per-individual basis using the SAMtools v1.18 *coverage* function (Danecek et al. 2021), and individuals with an average depth lower than 0.5X for any chromosome were subsequently removed from the data set prior to further analysis. We then generated a linkage disequilibrium (LD) pruned dataset by first subsetting one tenth of the data (by selecting every 10^th^ variant) and running the ngsLD program (v1.2.0), which estimates pairwise LD by considering the genotype uncertainty involved with using genotype likelihoods (Fox et al. 2019) to generate a list of unlinked variants. We then ran ANGSD again to output final genotype likelihoods for only these unlinked variants.

### Bioinformatic Processing of mtDNA Data

Bioinformatic processing of raw mtDNA data followed the same procedures described above for the nuclear data up to and including the indel realignment step. We then followed the procedures described in Therkildsen & Palumbi, (2017) to call mtDNA variants using freebayes v1.2.0 (Garrison and Marth 2012) in haploid mode, requiring at least 3 reads per sample for an alternate allele to be considered (-min-alternate-count 3), and with a minimum base quality and mapping quality of 30. Multi-allelic sites were then removed, and the most common alternate allele picked using the vcflib package and vcfflatten function (Garrison et al. 2022).

### Admixture Analyses

To investigate the potential presence of *A. anguilla* admixture in our Newfoundland eel samples, we conducted a principal components analysis (PCA) using the PCAngsd (v1.21) program (Meisner and Albrechtsen 2018) using the full SNP dataset containing both *A. anguilla* and *A. rostrata*. Similarly, we also performed a PCA on the mitogenome variant data using the - pca function in plink v1.90 (Purcell et al., 2007). Finally, we used the program NGSadmix (v23, Skotte et al., (2013)) on the LD-pruned nuclear variant dataset (see above) to assess levels of European admixture in our Newfoundland eel samples.

## Results

### Environmental DNA Survey

*Anguilla anguilla* mtDNA was detected with eDNA metabarcoding at eight sites in southern Newfoundland (Figure 2a), at three of these sites with the FISHE marker and at the remaining five with the MIFISHU marker, in a single sample at each site (Figure 2b). In contrast, *A. rostrata* were detected at the majority of sites sampled around the island, including at the eight sites where *A. anguilla* were detected, and were detected in multiple samples per site (Figures 2a and 2b). Two unique amplicon sequence variants (ASVs) assigned to *A. anguilla* for the FISHE marker with >=99% sequence similarity score (percent identity x percent query coverage, as output by BLAST results), while a single ASV assigned to *A. anguilla* for the MIFISHU marker. Sequencing depths (after correction for potential contamination) for four out of five MIFISHU *A. anguilla* detections fell within the 25^th^ (n=3; sites HIR, ROB, and SLM) and 50^th^ (n=2; sites BIR and DBB) percentiles for sequencing depths of all fish species detected at those sites, with the detection at site ROB having a very low absolute depth (Figure 2b). Sequencing depths for the three sites with FISHE *A. anguilla* detections (both ASVs pooled) fell within the 25^th^ percentile for sequencing depths of all species detected at those sites (Figure 2b).

**Figure 2.**
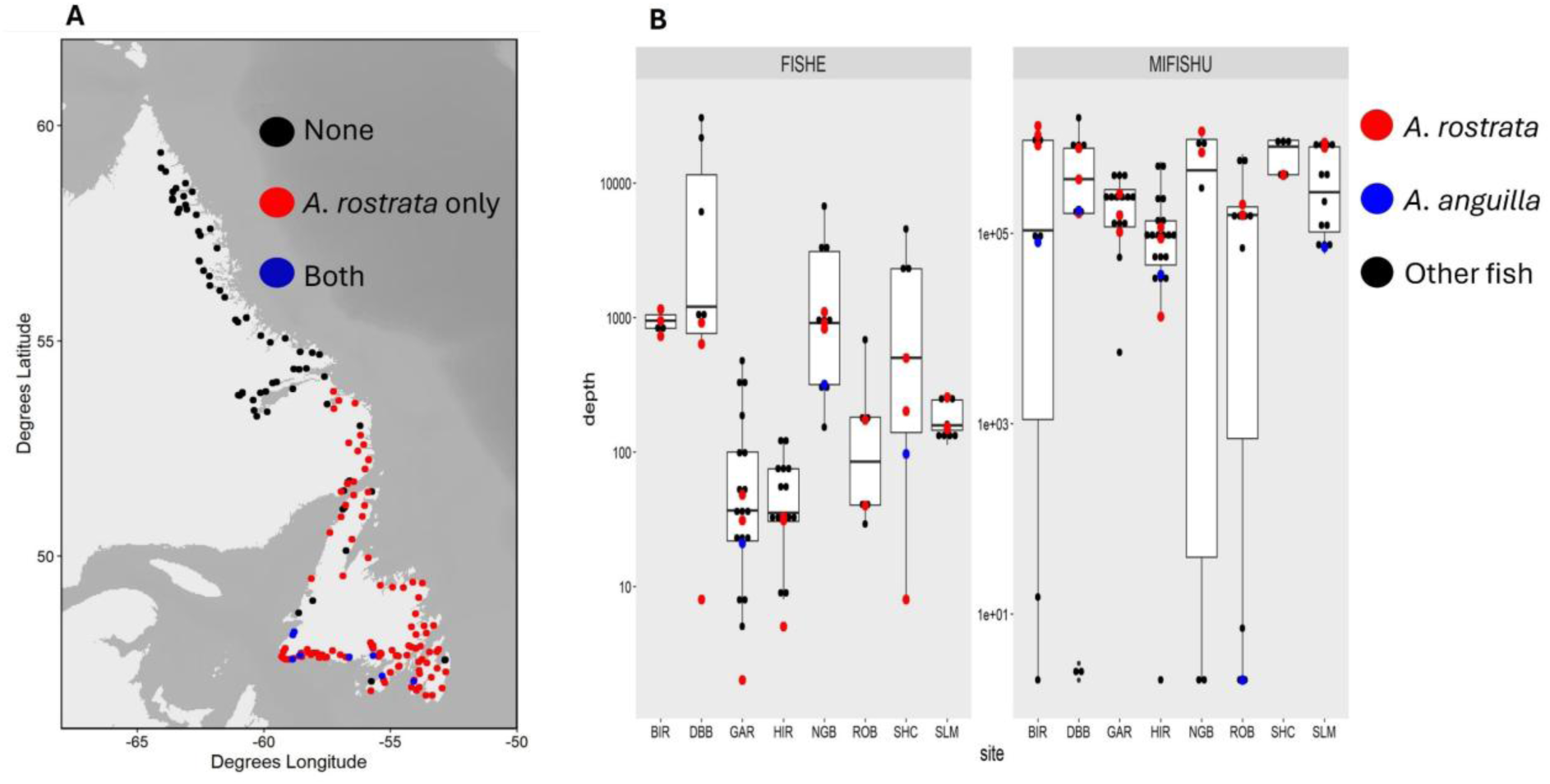
A) Locations of sampling with eDNA metabarcoding in 2019-2021, showing sites where no eel DNA was detected (black points), sites where only *A. rostrata* DNA was detected (red points), and sites where both *A. rostrata* and *A. anguilla* DNA were detected (blue points); B) Sequencing depths (i.e., read counts), across samples for each site where *A. anguilla* DNA was detected, for ASVs assigning to *A. anguilla* (blue points), *A. rostrata* (red points), and other fish species (black points) for the FISHE and MIFISHU markers. Note the log scale on the y-axis. Boxplot boxes represent the interquartile range (IQR; distance between 25th and 75th percentiles), while upper and lower whiskers extend to the largest and smallest values (respectively) that are no further than 1.5x the IQR.

### Mitochondrial Genotyping

After removal of 27 individuals with <15x coverage of their mitochondrial genomes, we genotyped a total of 813 individuals’ mitochondrial genomes (Newfoundland samples and those from Enbody et al., (2021)) and found a total of 2887 variants (2878 SNPs) out of the total 16,683 positions in the *A. anguilla* reference mitogenome. A principal components analysis on the mtDNA variants explained 46.3% and 4.4% of the variance in the data the first two components (respectively; for a PCA with a total of 20 components) and showed clear clustering of the two species along the first component axis (Figure 3a). The exceptions to this clustering were three individuals sampled at Ship Cove, Newfoundland, which cluster with the *A. anguilla* samples and thus appear to have mitogenomes of European ancestry.

**Figure 3.**
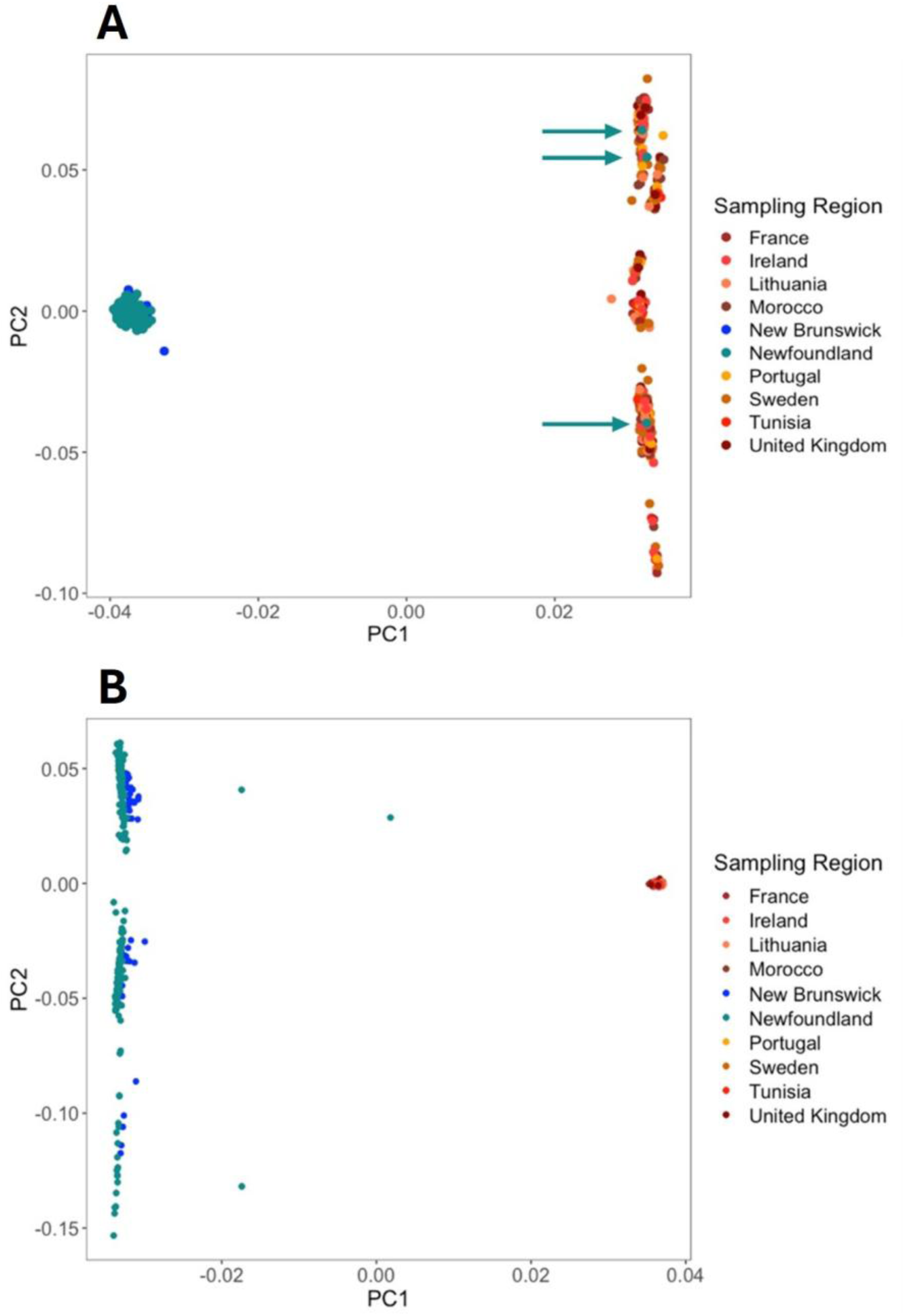
A) Biplot of the first two principal components from a PCA on mtDNA variants (as determined through Freebayes) for samples collected in NL as well as those from Enbody et al. 2021 (n=325 and n=488, respectively). PC1 explained 46.3% of the variance in the data, and PC2 explained 4.4% of the variance. Arrows indicate the three Newfoundland samples clustering with the European Eel samples. B) Biplot of first two principal components for a PCA on nuclear SNPs across 19 chromosomes for samples from NL and Enbody et al. (2021) (n=327 and n=488, respectively). PC1 explained 16.6% of the variance in the data, and PC2 explained 0.16%.

### Nuclear Genotyping

Generation of genotype likelihoods for 840 samples (Newfoundland samples and those from Enbody et al., (2021)) resulted in 25,338,859 SNPs across the 19 chromosomes. Twenty-five samples were subsequently removed and excluded from further analysis due to very low mean depth (< 0.5X) at a minimum of one chromosome. A PCA done on all nuclear SNPs explained 16.6% and 0.16% of the variance in the data in the first two components (respectively) and revealed marked clustering of species along the first axis with the exception of the same three individuals from Ship Cove, which fall intermediate to the main species clusters along PC1 (Figure 3b). Admixture results on the LD-pruned SNPs (n=1,938,332) show very low levels of *A. anguilla* admixture in samples from North America, and vice versa (Figure 4a), with once again the exception being the three samples from Ship Cove, which had European Eel admixture proportions of 0.224, 0.217, and 0.486, characteristic of two backcrossed individuals (American Eel direction) and one first-generation (F1) hybrid (Figure 4b).

**Figure 4.**
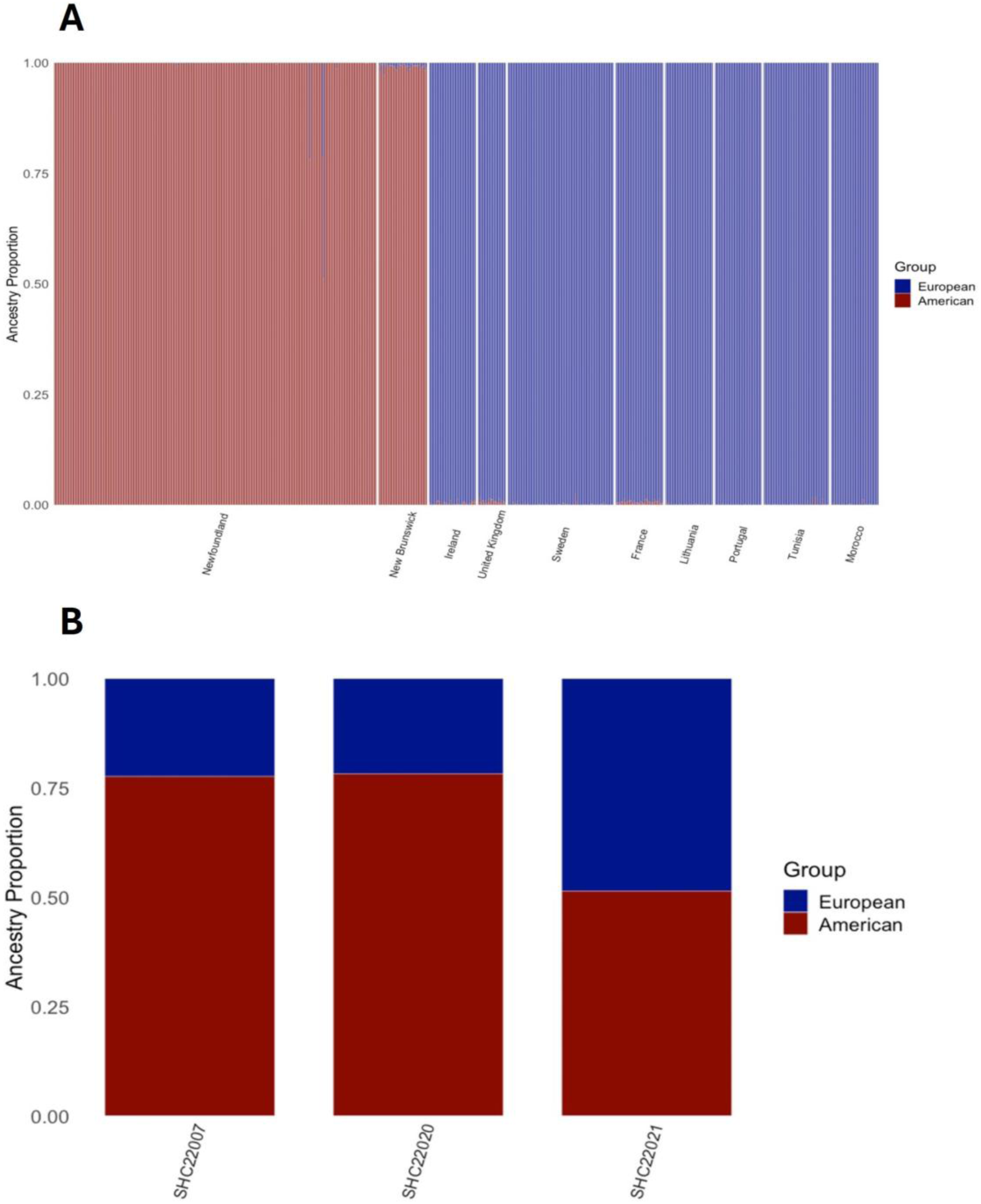
A) Admixture proportions of European vs. American Eel ancestry in all samples collected in NL and from Enbody et al. 2021, as generated through ngsadmix; and B) enlarged view of admixture proportions of European vs. American ancestry in three putative hybrids samples collected from Ship Cove, Newfoundland.

## Discussion

Previous detection of American-European Eel hybrids (Avise et al. 1990; Albert et al. 2006; Gagnaire et al. 2009) demonstrates that hybridization between the species does occur; however, evidence of hybrids has been geographically restricted. Our results suggest that American-European Eel hybrids occur outside of the previously known zone of hybrid occurrence (Iceland), with detections of European Eel mtDNA in an eDNA survey at eight sites in southern Newfoundland, and whole genome sequencing of eels collected at sites in this region showing the presence of one putative first generation hybrid and two backcrossed individuals (all with *A. anguilla* mitogenomes) at one of these sites two years later. These detections suggest that the larval dispersal and isolation of these two species may be more nuanced than originally thought, prompting further examination of hybridization in particular.

Our eDNA analyses extend the results of Crowley et al., (2024) and suggest the unexpected presence of *A. anguilla* is largely geographically restricted to rivers along the south coast of Newfoundland. While eDNA can have limitations in relation to false positives due to contamination or reference database issues (reviewed by Miya 2022), several factors provided us with confidence to further investigate our *A. anguilla* ASV detections as a true signal of this species’ presence. These detections fell within a fairly localized region (rather than at random within the survey area), had relatively high amplicon sequence variant signal strength (read counts) after stringent correction for potential contamination, and were detected by two different mtDNA markers. Indeed, this scenario perfectly illustrates eDNA’s effectiveness as an early detection method for unexpected species’ occurrences (i.e. Larson et al. 2020), and the results we report here demonstrate the complementarity of eDNA with more targeted sampling methods to form and address new research questions.

The whole genome sequencing data provide further support for the occurrence of eels with European ancestry in southern Newfoundland. The mitochondrial whole genome data showed three individuals (all at the Ship Cove site) had European-type mitogenomes, while the nuclear data showed these three fish to be hybrids. The maternal inheritance of mtDNA limits the ability of the eDNA survey to detect hybridization to one direction in this region (i.e. individuals from a mother with a European-type mitogenome); however, the combination of nuclear and mitochondrial data analyses here show that for the three hybrids detected, hybridization is indeed unidirectional. While we cannot rule out from our small sample size that hybridization does not occur in the opposite direction in the region as well, nonetheless these results are in line with some previous work observing asymmetric (American to European) patterns of admixture (Gagnaire et al. 2009; Wielgoss et al. 2014).

Another interesting observation in the nuclear data is the three clusters of *A. rostrata* individuals in the PCA biplot. These likely reflect the presence of putative structural variants reported by Ulmo-Diaz et al., (2023) on chromosomes 11 and 13 (and indeed, we do see similar clustering patterns on PCAs of these two chromosomes individually; see Supplementary Figures). It is intriguing that the putative F1 hybrid and one of the putative backcrossed individuals fall within the same cluster (i.e. homozygous for one version of the variant), while the other backcrossed individual falls within the other homozygote cluster (this is clearly observed in the chromosome 11 PCA as well, while results for chromosome 13 are less clear). Future work investigating the frequency of these potential structural variants in pure and hybrid individuals could bring interesting insight to the nature of hybridization between these two species.

A few key questions arise from our results, to which we can only speculate answers. The first and most fundamental question: is the presence of hybrid eels outside of Iceland restricted to southern Newfoundland? While our targeted sampling was limited to 15 sites, the eDNA survey was extensive, with over 100 sites in Newfoundland in 2020, and an additional 67 sites in Labrador between 2019 and 2021 (see Crowley et al. 2024). Given that eDNA has proven to be a very sensitive sampling method (e.g. Boivin-Delisle et al. 2021; McColl-Gausden et al. 2021), and that methods were consistent across the survey in this study, it is reasonable to assume that the detections of *A. anguilla* eDNA provide a plausible picture of a localized rather than widespread pattern of hybrid occurrence in the province of Newfoundland and Labrador during the years sampled.

Beyond Newfoundland and Labrador, the potential for hybrid occurrence in the continental Americas is more difficult to formally assess. Prior to this study, the vast majority of observations of hybrids of any class have been geographically restricted to Iceland, and observed mitochondrial genome lineages of the two species have remained distinct outside of Iceland (i.e. Wielgoss et al. 2014). Nonetheless, given that past work has reported limited levels of *A. anguilla* admixture in eels sampled in North America (e.g. Albert et al. 2006; Wielgoss et al. 2014), hybrids may be present outside of Newfoundland with more frequency than previously thought, and may have simply gone undetected. Additionally, given that *A. anguilla* is an unexpected species in the continental Americas, previous eDNA surveys that discarded unexpected taxa or used custom reference databases with only known or native species would not have reported hybrid detections. Choice of reference database and reliance on prior knowledge of species’ distributions should be carefully considered when conducting eDNA work, as these can contribute to confirmation bias and ultimately lead to missing or screening out unexpected species when they are actually present. This is especially critical in environments that may be undergoing changes in species distributions due to climate and/or human activities, and evidence such as number of markers showing detection, spatial distribution of detections, and relative distribution of read depths should be evaluated for unexpected detections before simply excluding them based on prior expectations. Future work on these two species could leverage eDNA surveys to identify sites of potential hybrid occurrence, followed by more targeted sampling to confirm hybrid presence, as we show here. Additionally, previous eDNA metabarcoding datasets from the Americas could be revisited to screen for *A. anguilla* detections that may have been incidentally deemed erroneous and subsequently removed. Ultimately, knowledge of the extent of hybrid occurrence in North America will be required in order to effectively investigate the mechanisms that drive these patterns of hybrid occurrence.

This leads to the second key question: what mechanisms drive hybrid dispersal to southern Newfoundland? The species are thought to spawn in separate but overlapping areas characterized by thermal fronts in the Subtropical Convergence Zone (STCZ) in the southern Sargasso Sea, with the centre of *A. rostrata* spawning more westerly than that of *A. anguilla* (e.g. Kleckner et al. 1983; Kleckner and Mccleave’ 1988; Munk et al. 2010). Initial hypotheses proposed that larvae of both species drift west, then north via entrainment in the Antilles and Florida Currents, the Gulf Stream and North Atlantic Drift. Segregation to different continents is thought to be due to *A. rostrata*’s shorter larval stage duration and thus earlier exit from the main currents (Bonhommeau et al., 2009; Kettle & Haines, 2006; Power & McCleave, 1983; Schmidt, 1923). Differences in gene expression timing in a wide range of biological functions between larvae of the two species may influence this developmental timing and therefore dispersal patterns (Bernatchez et al. 2011). While an alternative larval drift route for *A. anguilla* (the eastward moving Subtropical Counter Current) has also been proposed (Miller et al. 2009; Munk et al. 2010, 2023), if hybrids have a larval stage duration and/or route that is intermediate to those of the pure species, they would be expected to metamorphosize into glass eels at a location intermediate to the Americas and Europe, such as Iceland (Avise et al. 1990, Albert et al. 2006). However, Iceland is actually located at the western edge of the *A. anguilla* freshwater range rather than intermediate to the two species’ ranges, and indeed the majority of eels in Iceland have been found to be genetically pure *A. anguilla* (Albert et al. 2006). As such, perhaps this phenomenon occurs in the western Atlantic as well, with the intermediate larval attributes of hybrids interacting with environmental factors and resulting in some hybrids dispersing to the eastern-most part of the *A. rostrata* range (i.e. Newfoundland).

Finally, are hybrids in southern Newfoundland a new phenomenon, or an old but previously undetected one? Temperature in the Sargasso has been increasing since the 1970s, shifting the 22.5 degree C isotherm that delineates the northern limit of the eel spawning range (Friedland et al. 2007). Perhaps, a shift in the two species’ spawning ranges could alter the direction in which some hybrid larvae are advected, leading to their more frequent occurrence outside of Iceland. Without knowing the ages of the hybrids in our study, it is not possible to assess whether they represent a single or multiple dispersal events. Continued monitoring (both with eDNA and more traditional sampling methods) of sites in the region will be required to validate whether the patterns of hybrid occurrence we report here are stable both spatially and temporally.

To conclude, our results show that first-generation and backcross American-European hybrid eels can be found in southern Newfoundland, Canada. These findings represent the first observations of hybrid eels in Newfoundland and Labrador, the first validated observation of an F1 hybrid outside of Iceland, and the first observation of European mitochondrial genomes in eels collected in the Americas, to our knowledge. This work not only invites re-examination of previous hypotheses regarding Atlantic eel hybridization and larval hybrid dispersal, but also demonstrates the utility of pairing eDNA surveys as an early detection tool for non-native species with subsequent targeted sampling to obtain more detailed information regarding the specifics of hybridization. This exciting discovery opens the door for more work investigating the potential for the presence of hybrid eels elsewhere in the two species’ continental ranges, the mechanisms that determine these patterns of distribution, and whether or not they are changing over time. For eels, whose unique life history attributes remain poorly understood, with each piece of the puzzle uncovered we become better equipped to manage and conserve them.

## Acknowledgements

Authors would like to acknowledge funding sources from the Killam Doctoral Award (SC), Fisheries and Oceans Canada Genomics Research and Development Initiative, and NSERC Discovery Grants (IRB and PB). Authors also acknowledge staff of Fisheries and Oceans Canada in the Newfoundland and Labrador Salmonids Section for assistance with eDNA and targeted eel sampling, as well as Canadian Coast Guard helicopter pilots for assistance with accessing eDNA sampling sites. For processing of eDNA samples, we thank Hoda Rajabi, Emily Porter, Eman Elbakry, Jasmine Yeung, Gary McKeown, Patrick Banks, Muneesh Kaushal, Gregory Singer, and Beverly McClenaghan of the Centre for Environmental Genomics Applications.

