## Supplementary Figures for "eDNA metabarcoding and whole genome sequencing detect European-American Eel hybrids in northeastern Canada"

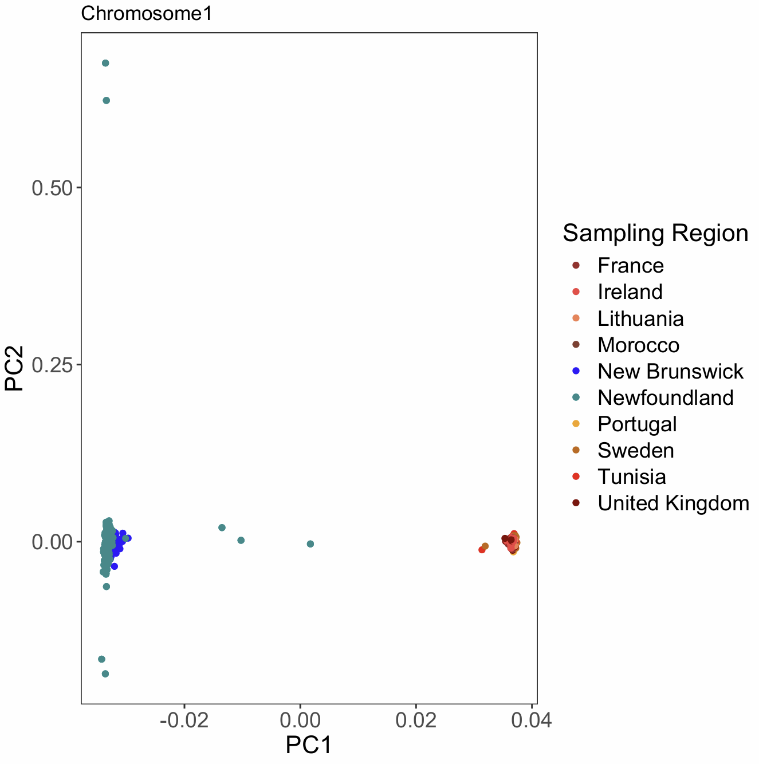

Supplemental Figure 1. Biplot of first two principal components for a PCA on nuclear SNPs across chromosome 1 for samples from NL and Enbody et al. (2021) (n=327 and n=488, respectively).

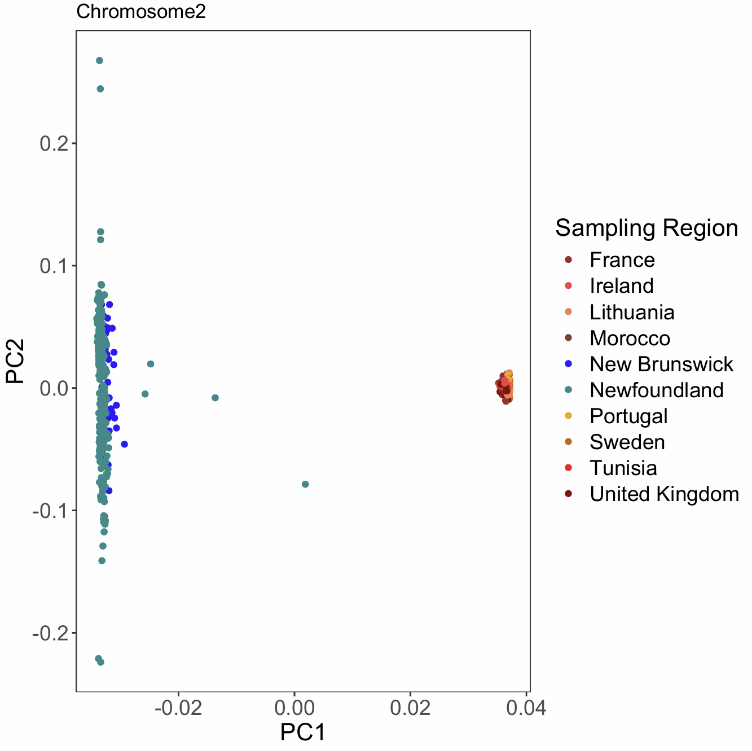

Supplemental Figure 2. Biplot of first two principal components for a PCA on nuclear SNPs across chromosome 2 for samples from NL and Enbody et al. (2021) (n=327 and n=488, respectively).

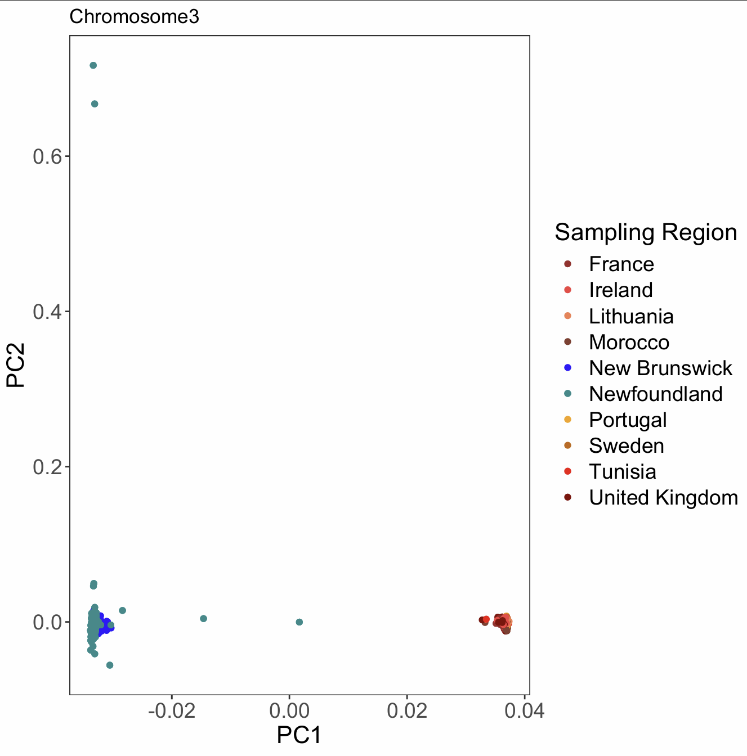

Supplemental Figure 3. Biplot of first two principal components for a PCA on nuclear SNPs across chromosome 3 for samples from NL and Enbody et al. (2021) (n=327 and n=488, respectively).

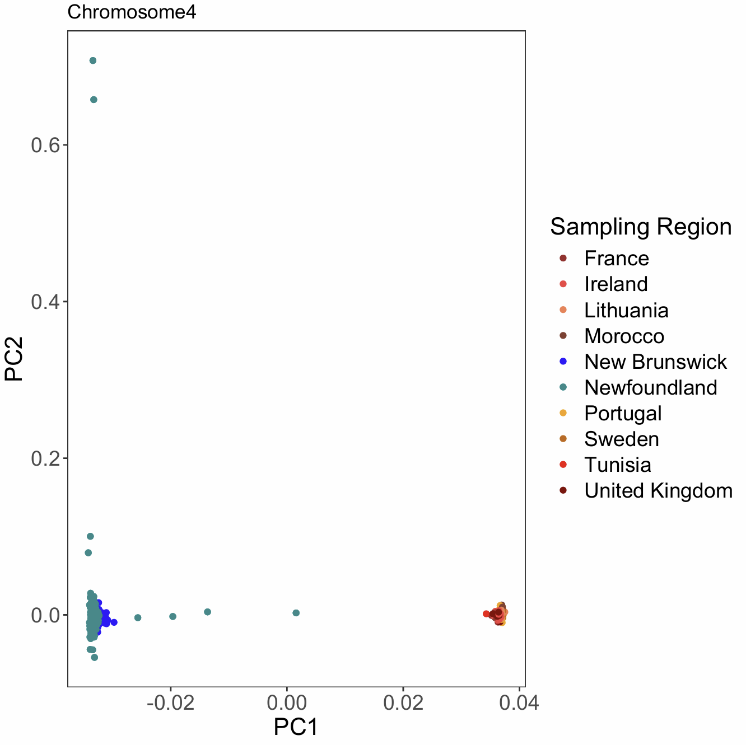

Supplemental Figure 4. Biplot of first two principal components for a PCA on nuclear SNPs across chromosome 4 for samples from NL and Enbody et al. (2021) (n=327 and n=488, respectively).

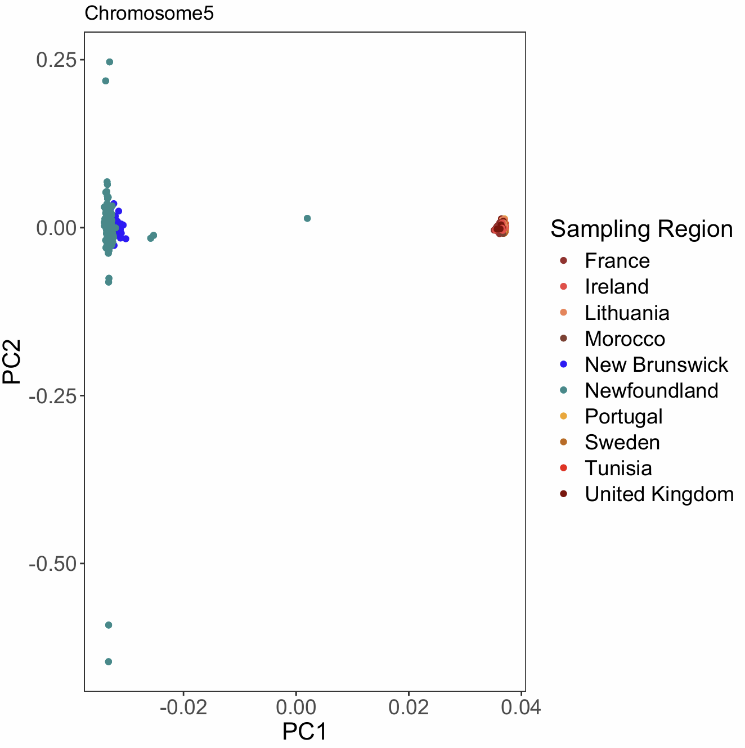

Supplemental Figure 5. Biplot of first two principal components for a PCA on nuclear SNPs across chromosome 5 for samples from NL and Enbody et al. (2021) (n=327 and n=488, respectively).

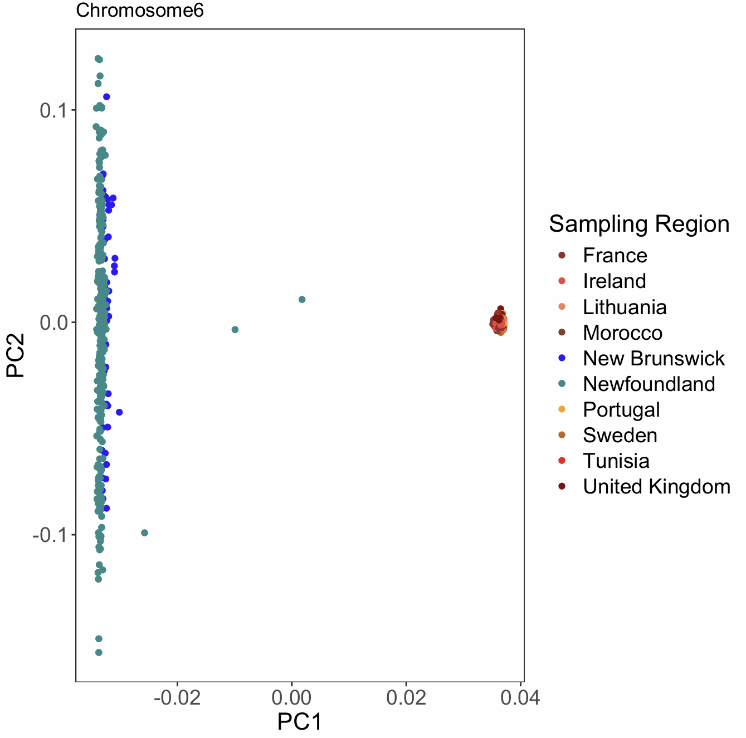

Supplemental Figure 6. Biplot of first two principal components for a PCA on nuclear SNPs across chromosome 6 for samples from NL and Enbody et al. (2021) (n=327 and n=488, respectively).

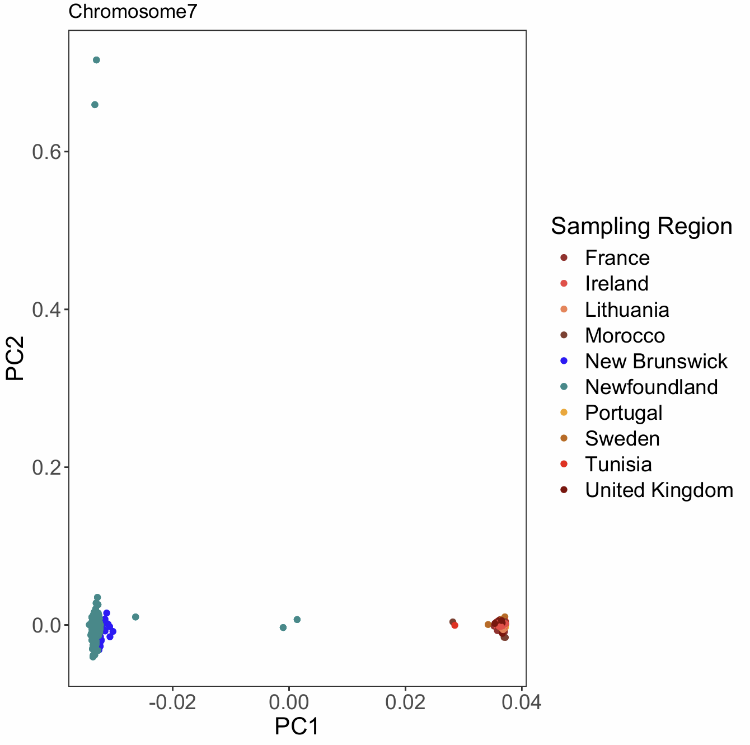

Supplemental Figure 7. Biplot of first two principal components for a PCA on nuclear SNPs across chromosome 7 for samples from NL and Enbody et al. (2021) (n=327 and n=488, respectively).

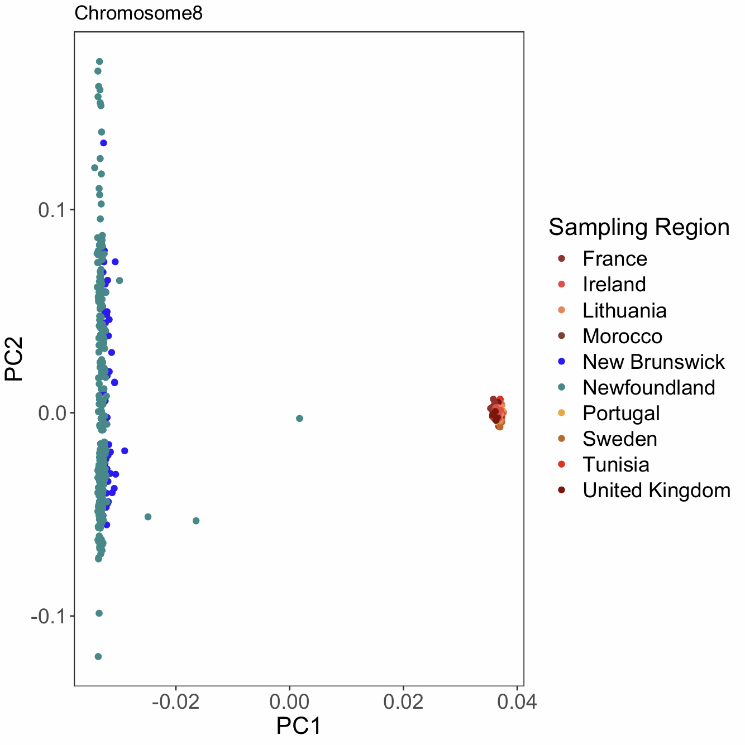

Supplemental Figure 8. Biplot of first two principal components for a PCA on nuclear SNPs across chromosome 8 for samples from NL and Enbody et al. (2021) (n=327 and n=488, respectively).

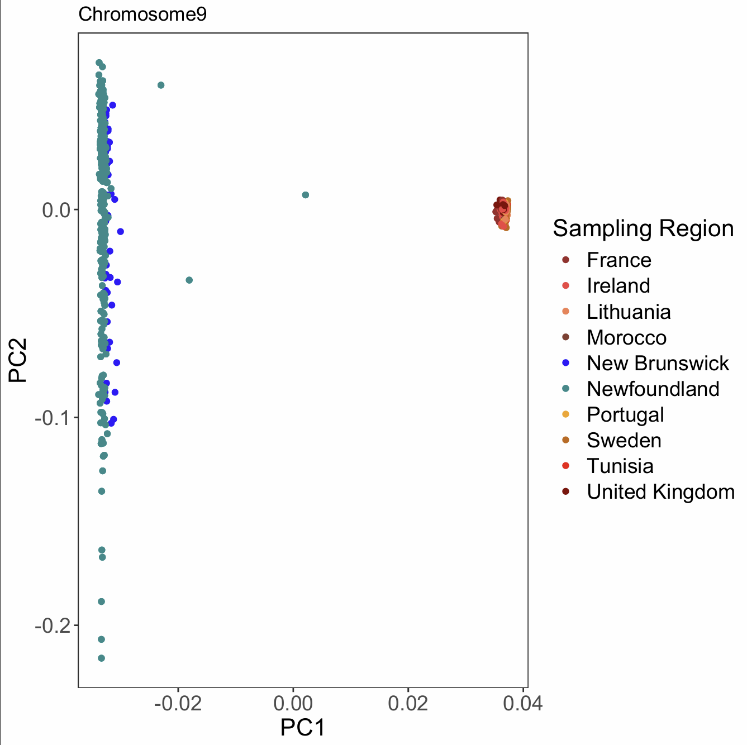

Supplemental Figure 9. Biplot of first two principal components for a PCA on nuclear SNPs across chromosome 9 for samples from NL and Enbody et al. (2021) (n=327 and n=488, respectively).

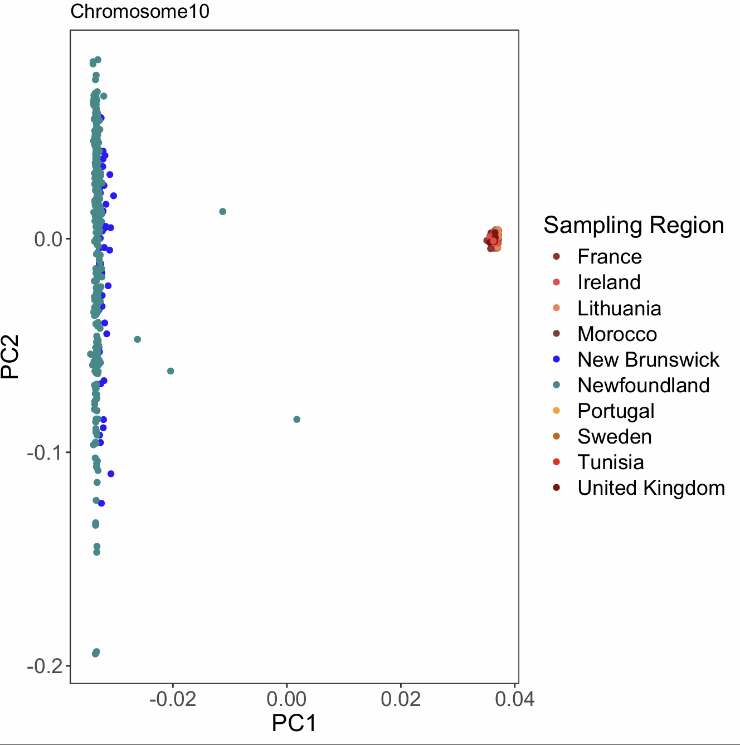

Supplemental Figure 10. Biplot of first two principal components for a PCA on nuclear SNPs across chromosome 10 for samples from NL and Enbody et al. (2021) (n=327 and n=488, respectively).

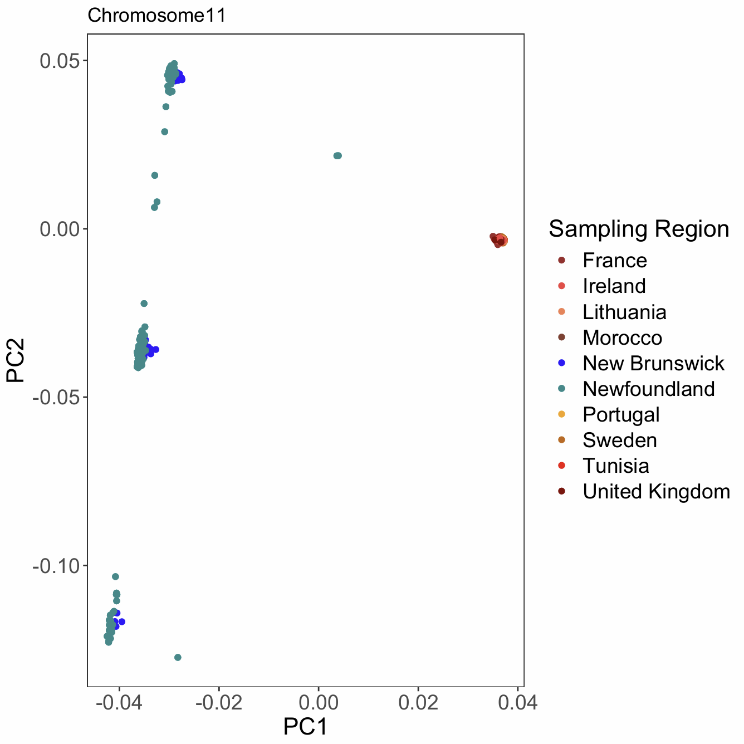

Supplemental Figure 11. Biplot of first two principal components for a PCA on nuclear SNPs across chromosome 11 for samples from NL and Enbody et al. (2021) (n=327 and n=488, respectively).

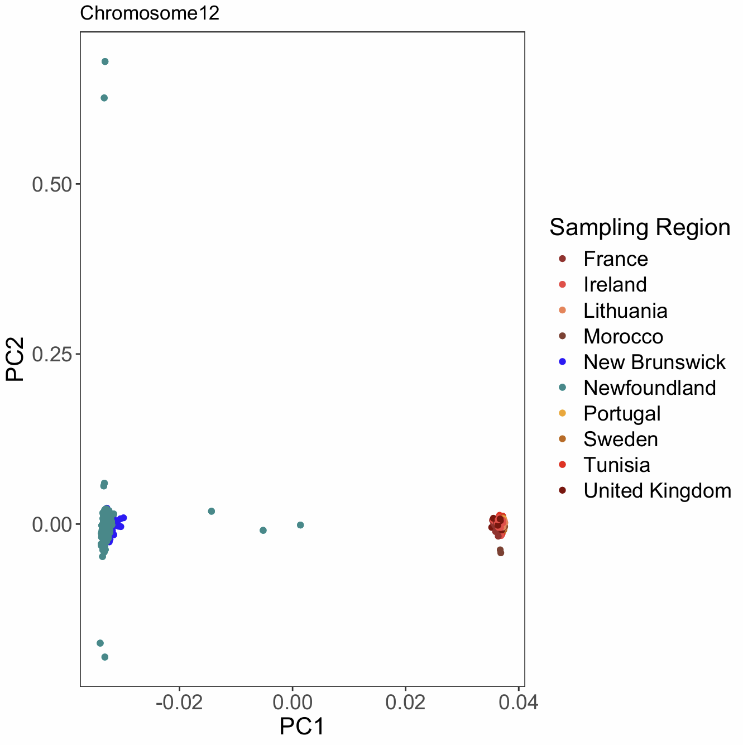

Supplemental Figure 12. Biplot of first two principal components for a PCA on nuclear SNPs across chromosome 12 for samples from NL and Enbody et al. (2021) (n=327 and n=488, respectively).

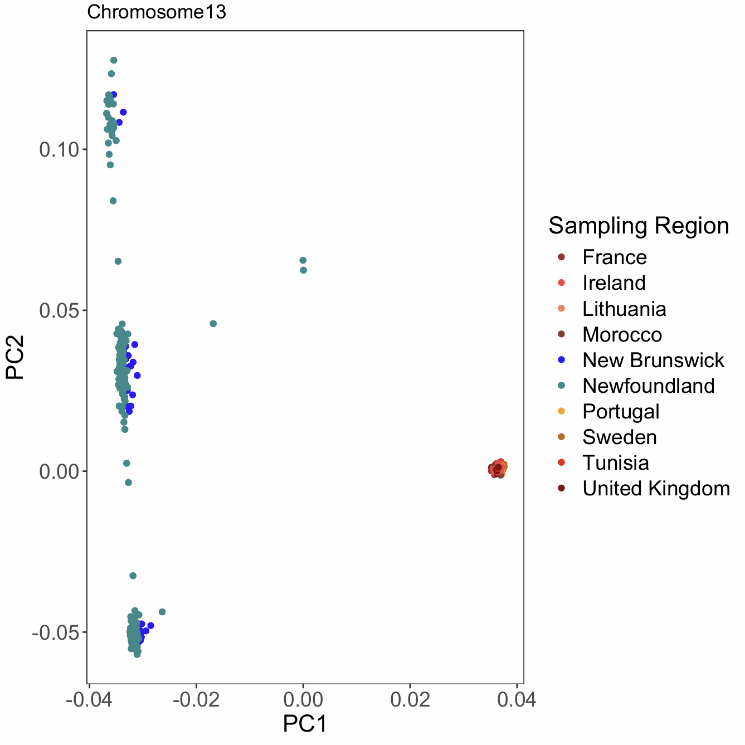

Supplemental Figure 13. Biplot of first two principal components for a PCA on nuclear SNPs across chromosome 13 for samples from NL and Enbody et al. (2021) (n=327 and n=488, respectively).

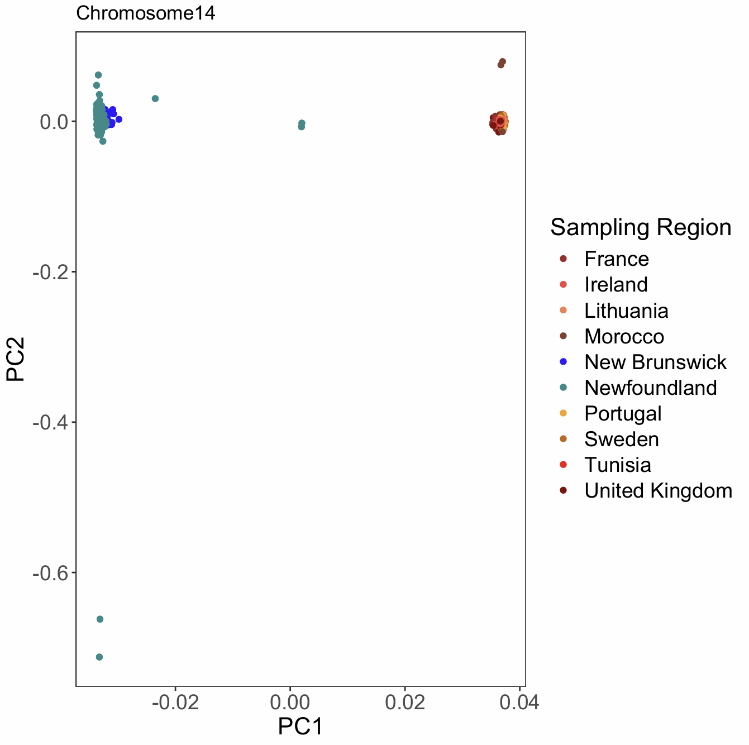

Supplemental Figure 14. Biplot of first two principal components for a PCA on nuclear SNPs across chromosome 14 for samples from NL and Enbody et al. (2021) (n=327 and n=488, respectively).

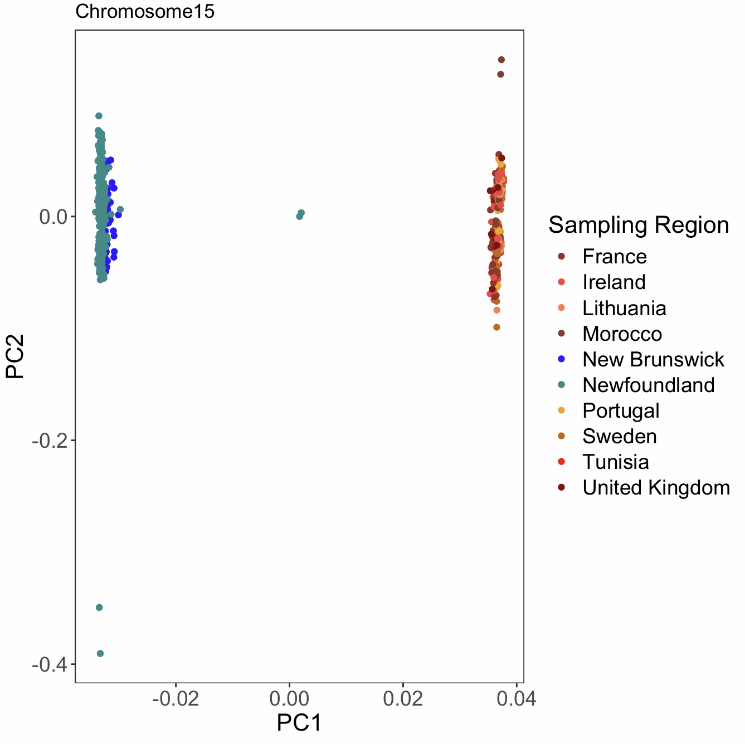

Supplemental Figure 15. Biplot of first two principal components for a PCA on nuclear SNPs across chromosome 15 for samples from NL and Enbody et al. (2021) (n=327 and n=488, respectively).

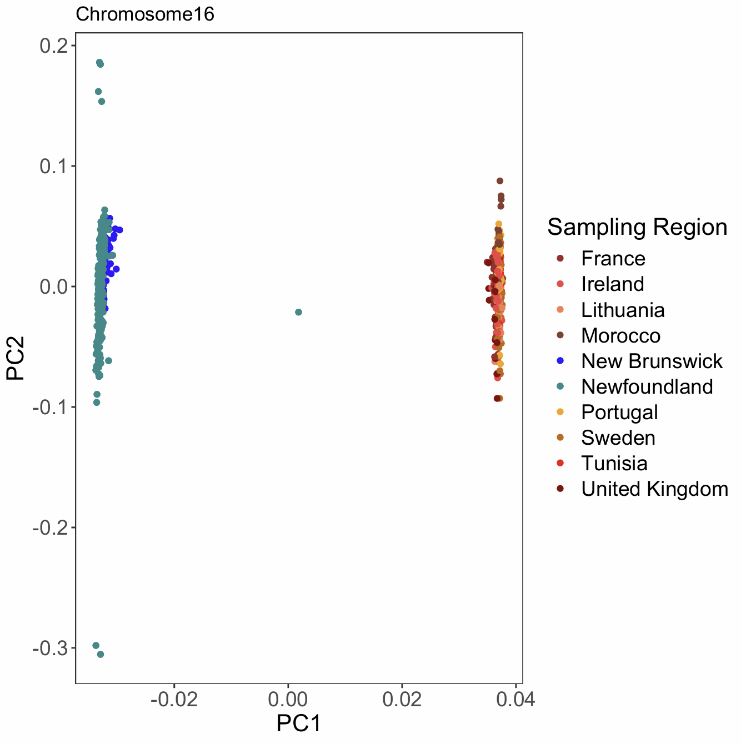

Supplemental Figure 16. Biplot of first two principal components for a PCA on nuclear SNPs across chromosome 16 for samples from NL and Enbody et al. (2021) (n=327 and n=488, respectively).

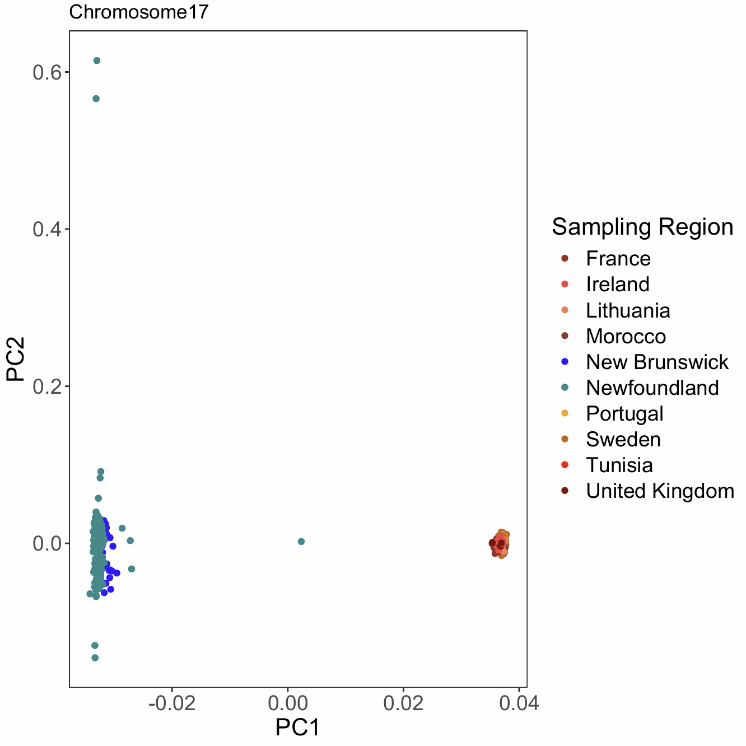

Supplemental Figure 17. Biplot of first two principal components for a PCA on nuclear SNPs across chromosome 17 for samples from NL and Enbody et al. (2021) (n=327 and n=488, respectively).

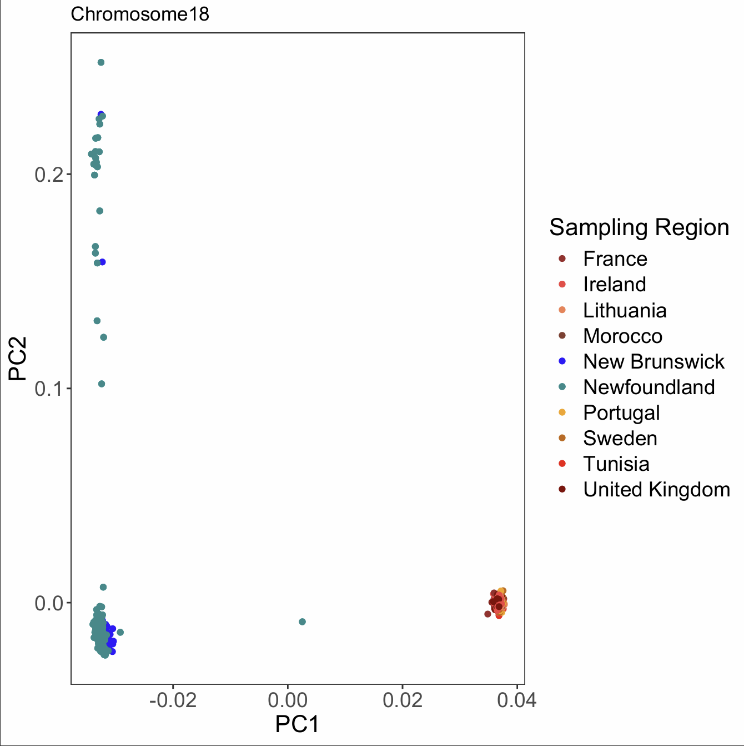

Supplemental Figure 18. Biplot of first two principal components for a PCA on nuclear SNPs across chromosome 18 for samples from NL and Enbody et al. (2021) (n=327 and n=488, respectively).

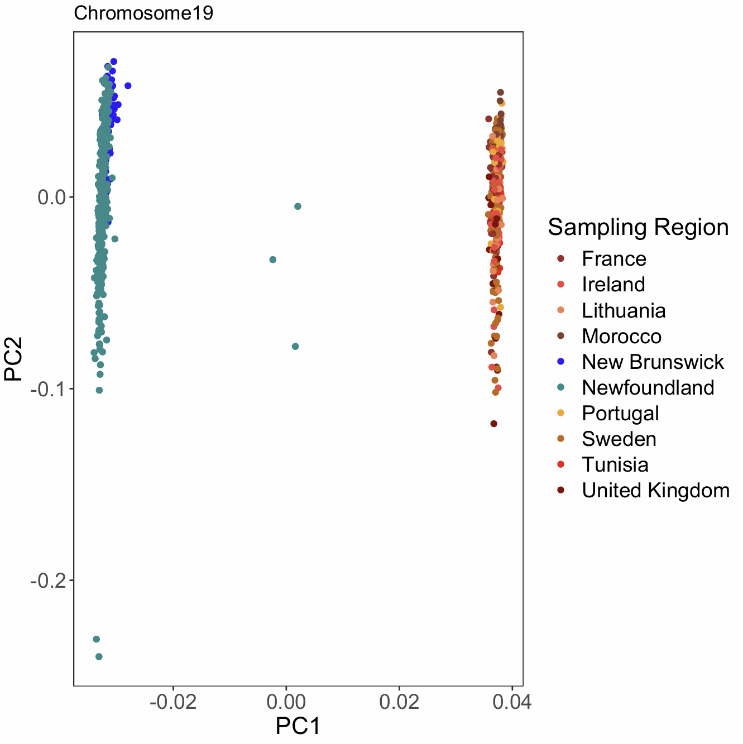

Supplemental Figure 19. Biplot of first two principal components for a PCA on nuclear SNPs across chromosome 19 for samples from NL and Enbody et al. (2021) (n=327 and n=488, respectively).
